# Disrupting chemotaxis stimulates adhesion by recruiting a putative c-di-GMP effector to the *Caulobacter crescentus* cell pole

**DOI:** 10.64898/2026.08.21.746235

**Authors:** Rachel I Salemi, David M Hershey

**Affiliations:** Department of Bacteriology, University of Wisconsin – Madison, Madison, WI, USA

## Abstract

Contact with solid surfaces activates signaling pathways that promote biofilm formation in many bacteria. The alphaproteobacterium *Caulobacter crescentus* uses its flagellum to sense surfaces and responds by synthesizing an adhesive called the holdfast. The *C. crescentus* surface sensing pathway can be activated by mutating genes required for the assembly of the flagellum or genes required for chemotaxis. However, flagellar assembly and chemotaxis mutations activate distinct surface sensing pathways that differ in the activation of the diguanylate cyclase PleD. Here, we used a genome-wide screen to identify *cmrA* (CCNA_02061) as a crucial determinant of hyperadhesion in the chemotaxis mutant Δ*cheYII*. Genetic analysis showed that *cmrA* is important for activation of PleD in a context-specific manner. It is dispensable in wild-type and late-stage flagellar (Δ*flgH*) mutant backgrounds but promotes adhesion in early-stage flagellar assembly (Δ*fliF*), chemotaxis (Δ*cheYII*) and stator (Δ*motB*) mutant backgrounds. Fluorescently tagged CmrA displays a mostly cytoplasmic localization in genetic backgrounds where *cmrA* is dispensable for adhesion but localizes to the cell pole in backgrounds where it regulates adhesion. Structural modeling indicates that CmrA is a degenerate, catalytically inactive GGDEF/EAL domain containing protein, but *cmrA* alleles with mutated conserved c-di-GMP coordinating residues are unable to support hyperadhesion. Our results indicate that altering the directional switching of MotAB stators recruits CmrA to the cell pole where it activates PleD to drive surface adaptation. Ultimately, this work underscores the complexity of flagellar surface sensing by highlighting how the many rotational states of the motor stimulate distinct but overlapping c-di-GMP signaling pathways

**Importance:** Bacteria often transition from a free-swimming state to form surface-attached communities called biofilms. The flagellum allows bacteria to sense surface contact and activate biofilm formation, yet how distinct structural states of this complex machine trigger surface sensing remains poorly understood. In this study, we identify CmrA as a key signaling link that senses flagellar motor disruption and activates second-messenger signaling to promote cell adhesion in *Caulobacter crescentus*. Our findings demonstrate that bacterial surface sensing is not a simple binary switch. Instead, distinct mechanical perturbations to the flagellum engage specialized, overlapping signaling pathways to fine-tune surface adaptation. Understanding these nuanced pathways will inform strategies to manipulate biofilm formation for human benefit.

## Introduction

Bacteria form multi-cellular structures called biofilms that contain sessile cells protected by an extracellular matrix (1, 2). Contact with solid surfaces often promotes biofilm formation by triggering surface sensing pathways (3). Surfaces are sensed mechanically by extracellular appendages such as flagella and type IV pili (4, 5). These appendages are best known for promoting cellular motility, but their involvement in surface recognition has also been well-documented (6, 7). Current models suggest that obstructions to type IV pili retraction or flagellar rotation during surface contact activate signaling pathways that promote attachment (6–9).

The flagellum contains a rotating extracellular filament that propels cells through liquid environments (10, 11). Assembly of the flagellum proceeds in a defined sequence (12). Hook basal body (HBB) formation begins with the assembly of membrane spanning (MS) and cytosolic (C) rings followed by the insertion of a transenvelope structure containing a rod, the peptidoglycan-ring (P-Ring), the lipopolysaccharide-ring (L-Ring) and an extracellular hook. The HBB secretes flagellin proteins that form a filament extending from the hook. Stator complexes that surround the C-ring use the proton motive force to rotate the HBB and its associated filament, generating force that propels the cell (13, 14). Effector proteins called CheYs can bind to the C-ring and cause the clockwise (CW) to counterclockwise (CCW) rotational bias of the flagellum to switch. Phosphorylation of CheYs in response to chemical cues influences directional switching of the motor in a way that allows bacteria to navigate their environment using a process known as chemotaxis (15). Both the flagellar assembly pathway and chemotaxis signaling pathway influence surface sensing (8, 9, 16–19), but how the conformational changes that occur during directional switching influence downstream signaling responses remains unclear.

Surface sensing systems appear to share a common signaling paradigm centered on cyclic dimeric GMP (c-di-GMP), a second messenger that suppresses motility and promotes sessile behaviors such as biofilm formation (20). Cellular c-di-GMP concentrations are determined by diguanylate cyclase (DGC) and phosphodiesterase (PDE) enzymes (21). DGCs synthesize c-di-GMP by condensing two GTP molecules via their characteristic GGDEF domain (22). PDEs hydrolyze c-di-GMP, with EAL domain enzymes producing pGpG and HD-GYP domain enzymes producing GMP (23). c-di-GMP promotes biofilm formation by binding to diverse effector proteins, some of which contain degenerate GGDEF and EAL domains that are incapable of catalysis but retain binding affinity for c-di-GMP (24). c-di-GMP effector proteins can be challenging to identify bioinformatically as there is no universal consensus sequence for c-di-GMP binding.

*Caulobacter crescentus* is an aquatic alphaproteobacterium that serves as a model for studying flagellar surface sensing (9). *C. crescentus* undergoes a distinct, dimorphic growth cycle. Motile swarmer cells contain a single flagellum at one cell pole and differentiate into stalked cells by shedding their flagella and extending the cell envelope to form a stalk (25). *C. crescentus* can secrete a polysaccharide called the holdfast during the swarmer-to-stalked cell transition that acts as an adhesin by anchoring stalked cells tightly to exogenous surfaces (26). Holdfast production is conditional, and contact with a surface is one of the environmental cues that stimulates holdfast production (3, 6, 7).

Mutating genes involved in flagellar assembly stimulates the *C. crescentus* surface sensing pathway, causing cells to increase holdfast production and become hyperadhesive (Figure 1A) (1, 8, 9). A genome wide screen in a flagellar mutant (Δ*flgH*) background identified two DGCs called PleD and DgcB as crucial actuators of hyperadhesion (1, 7, 9, 27). The contributions of these two enzymes in regulating holdfast production depends on the assembly status of the flagellum. Disrupting early stages of flagellar assembly by blocking the completion of the MS-and C-rings causes a hyperadhesive phenotype that is distinct from when late stages downstream of C-ring assembly are disrupted. In late-stage flagellar mutants, *pleD* and *dgcB* have additive roles in activating adhesion. Mutating either gene partially suppresses hyperadhesion, while simultaneous disruption of both DGCs makes cells completely non-adhesive. Activation of DgcB is thought to require functional MotAB stators, and *motB* mutants phenocopy *dgcB* mutants in late-stage flagellar mutants. In contrast, adhesion is entirely dependent on *pleD* in early-stage flagellar mutants. *dgcB* and *motAB* are thought to be dispensable in these mutants because the stators cannot functionally engage with the flagellar motor in the absence of the C-ring (9).

**Figure 1.**
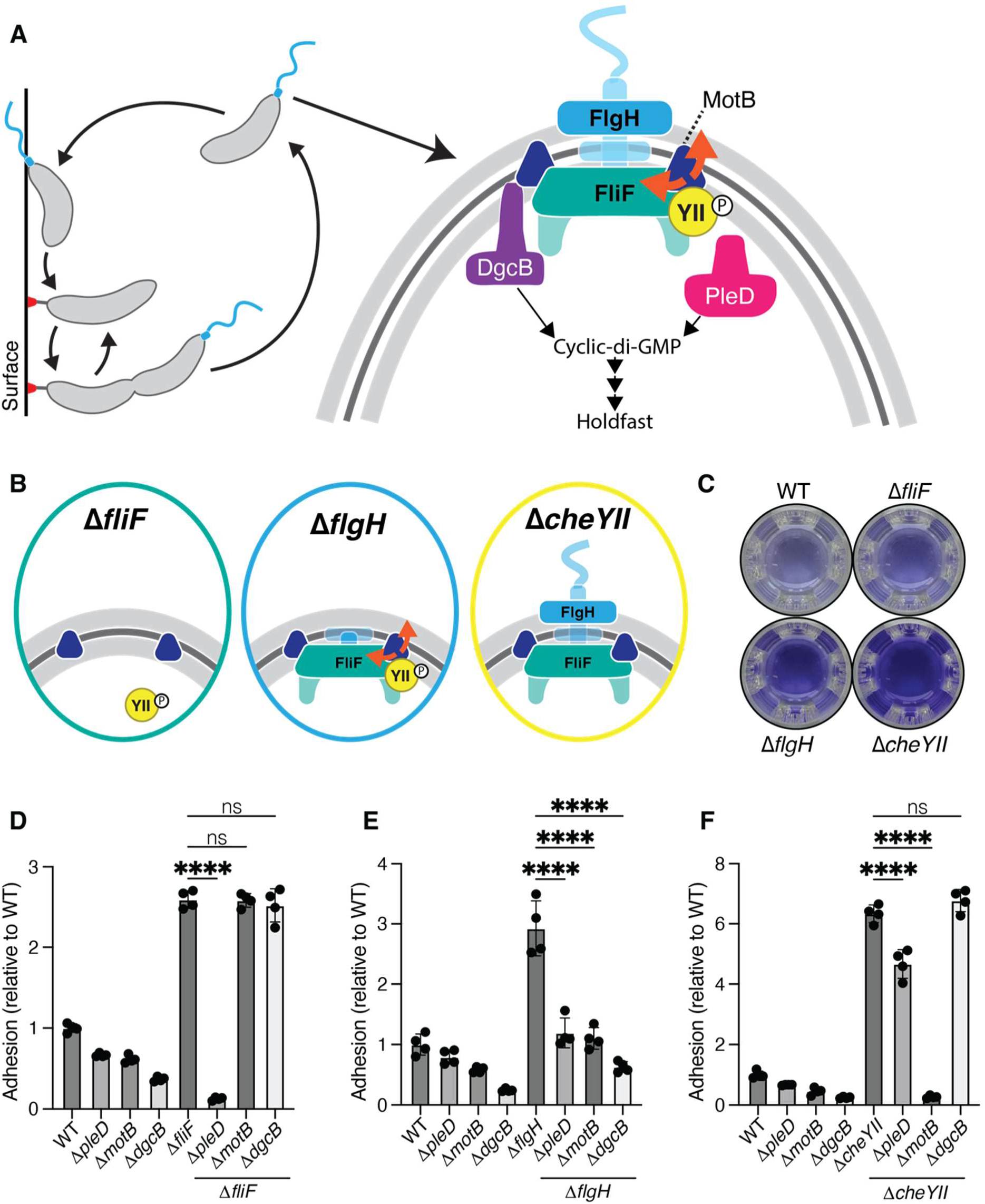
Disrupting the flagellum enhances adhesion. (A) Motile *Caulobacter crescentus* swarmer cells encounter a surface and respond by producing a holdfast polysaccharide. The flagellum is a multimeric protein complex that controls surface sensing. Disruption of this complex stimulates activity of the diguanylate cyclases PleD and DgcB, increasing cyclic-di-GMP concentrations and holdfast production. (B) Schematics of the flagellum in the absence of *fliF, flgH,* or *cheYII*. (C) Example of crystal violet staining in a 48-well plate in the WT, Δ*fliF,* Δ*flgH*, and Δ*cheYII* backgrounds. The darkness of the violet color indicates the efficacy of adhesion in each strain. (D-F) CV staining showing the relationship between *pleD, dgcB,* and *motB* adhesion in (D) Δ*fliF*, (E) Δ*flgH*, and (F) Δ*cheYII* backgrounds. Statistical significance determined by one-way ANOVA test and Tukey’s multiple-comparison test.

We recently showed that mutating genes required for chemotaxis also stimulates holdfast production without affecting flagellar assembly (16). Chemotaxis mutants with flagella that are locked in the CW rotational state display a hyperadhesive phenotype that differs from that of flagellar mutants. Mutating either *pleD* or *dgcB* in chemotaxis mutants has minimal effect on adhesion, but chemotaxis mutants with both DGCs disrupted are completely non-adhesive. The different DGC dependencies in the Δ*fliF* (MS-ring), Δ*flgH* (L-ring), and Δ*cheYII* (chemotaxis) backgrounds can be explained by interactions between the flagellum and the MotAB stator complex. In the absence of a flagellum (Δ*fliF*) the stators are unable to activate *dgcB* (9). In mutants that assemble the MS-ring and C-ring (Δ*flgH*), the stators are required for *dgcB* activation but do not affect *pleD* activation(9). When the flagellum is fully assembled (Δ*cheYII*) the stators activate both *pleD* and *dgcB* (16). This finding conflicts with previous assertions that the stators specifically activate DgcB and suggests the presence of a novel pathway for activating PleD.

In this study, we used a genetic screen to identify genes that promote *pleD* activation in the Δ*cheYII* background. We found that a previously uncharacterized gene (*CCNA_02061*), which we named <u>c</u>hemotaxis <u>m</u>odulated <u>r</u>epressor <u>A</u> (*cmrA*), is required for hyperadhesion when the flagellum is unable to switch its rotational direction. Fluorescently tagged CmrA exhibits a predominantly diffuse localization pattern in wild-type cells but localizes to the cell pole in mutant backgrounds in which directional switching of the flagellar motor is altered. A structural model predicts that CmrA contains a GGDEF domain and an EAL domain, both lacking the necessary residues for c-di-GMP metabolism. Mutating the GGDEF domain I-site or the predicted EAL c-di-GMP binding pocket abolishes the ability of *cmrA* to promote hyperadhesion. Together, our data support a model in which CmrA is recruited to the pole when chemotactic directional switching is suppressed and activates PleD.

## Results

Several mutations that disrupt flagellar function increase surface adhesion in *C. crescentus* (1, 8, 9, 16), but the signaling pathways vary depending on the mutant background. To formalize these signaling differences, we re-examined the genetic requirements for hyperadhesion in the Δ*fliF*, Δ*flgH*, and Δ*cheYII* mutant backgrounds (Fig. 1B). Crystal violet (CV) staining confirms that all three mutants exhibit a hyperadhesive phenotype (Fig. 1C). Consistent with previous results, each hyperadhesive mutant responds differently to the deletion of *pleD, dgcB,* or *motB* (Fig. 1D-F). Hyperadhesion in the Δ*fliF* mutant is dependent on *pleD,* while *dgcB* and *motB* are dispensable. Δ*cheYII* hyperadhesion requires *motB*, while *pleD* and *dgcB* are dispensable. In contrast to Δ*fliF* and Δ*cheYII* hyperadhesion, *pleD, dgcB,* and *motB* all have intermediate effects on Δ*flgH* hyperadhesion (9, 16). The requirement of *motB* for Δ*cheYII* hyperadhesion and the redundant relationship between *pleD* and *dgcB* contradicts the model that *pleD* is activated through a mechanism independent from *motB* (9, 16) and motivated our search for a gene that links MotAB function and PleD activation.

### *cmrA* is required for hyperadhesion in the Δ*cheYII* mutant background

Because *motB* is epistatic to *pleD* in the Δ*cheYII* background (16), we predicted the existence of one or more factors that activate *pleD* in a *motB*-dependent manner in chemotaxis mutants. These factors would not have been identified in the Δ*flgH* adhesion profiling experiment because *motB* does not contribute to *pleD* activation in late-stage flagellar mutants (9). We performed adhesion profiling (8) to identify genes required for adhesion in the Δ*cheYII* mutant. A barcoded transposon library (28) was developed in the Δ*cheYII* background and passaged in liquid medium with cheesecloth to enrich for non-adhesive mutants. We compared final passage fitness scores for the least adhesive transposon mutants in the Δ*cheYII* mutant to the least adhesive transposon mutants in the Δ*flgH* background (9) (Fig. 2A, Table S1). The results confirmed the epistasis patterns from Figure 1. Holdfast biosynthesis genes were required for adhesion in both the Δ*flgH* and Δ*cheYII* backgrounds. Disruption of *motB* impacted adhesion negatively in both backgrounds (Fig. 2A-B). Transposon insertions in *pleD* and *dgcB* were strong suppressors of hyperadhesion the Δ*flgH* background but did not affect adhesion in the Δ*cheYII* background. We also identified suites of genes required for hyperadhesion specifically in the Δ*cheYII* or Δ*flgH* backgrounds. Many previously reported *flagellar signaling suppressor* (*fss*) genes were represented in the Δ*flgH* specific suite (9, 16, 29). A separate set of genes affected adhesion specifically in the Δ*cheYII* background.

**Figure 2.**
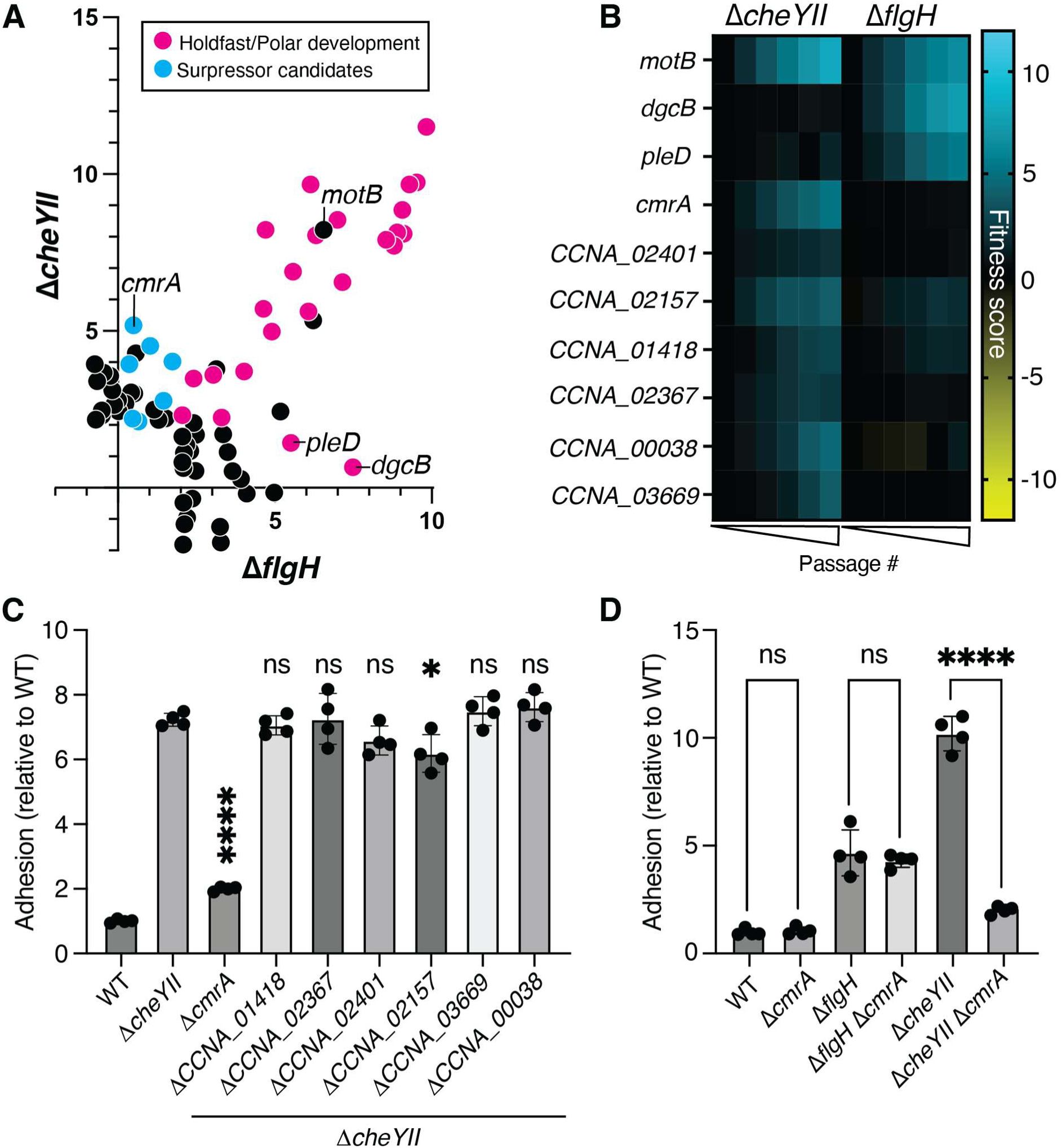
Identification of *cmrA* as a regulator of adhesion in Δ*cheYII* mutants (A) Fitness scores on the final day of passaging in the Δ*cheYII* background compared to the Δ*flgH* background identify adhesion control genes specific to each mutant. Each dot represents a different gene that has been interrupted by transposons. The pink dots represent genes that are involved in holdfast synthesis or polar development. Key genes are labeled (*cmrA, motB, pleD, dgcB*). (B) Heat maps showing the fitness scores of selected genes over the course of the 5 days of passaging. (C) CV staining of clean deletions of suppressor candidates in the Δ*cheYII* background. (D) CV Staining comparing *cmrA* suppression of adhesion in the wild-type, Δ*flgH*, and Δ*cheYII* backgrounds. For (C) and (D) Statistical significance was determined by one-way ANOVA test and Tukey’s multiple-comparison test. One asterisk indicates a P-value < 0.05 and four asterisks indicate a P-value < 0.0001.

We selected seven Δ*cheYII*-specific suppressor candidates for further study (Fig. 2B), generated in-frame deletions of each gene in the Δ*cheYII* background and measured adhesion using CV staining. While most of the mutations caused limited or no reversion of the hyperadhesive phenotype, mutating one gene (CCNA_02061) strongly suppressed Δ*cheYII*-dependent hyperadhesion. We named this gene *cmrA* (*<u>c</u>hemotaxis <u>m</u>odulated <u>r</u>epressor A*). An in-frame deletion of *cmrA* (Δ*cmrA*) had no effect on adhesion in a WT or Δ*flgH* background but strongly suppressed Δ*cheYII* hyperadhesion (Fig. 2D), confirming the specific role of *cmrA* in the chemotaxis mutant background.

### *cmrA* contributes to *pleD-*dependent hyperadhesion in specific motility mutants

To determine if *cmrA* is involved in activation of *pleD*, we performed epistasis analysis using CV staining to quantify adhesion (Fig. 3). Simultaneous inactivation of PleD and DgcB abolishes adhesion across all genetic backgrounds (Fig 1). Thus, if *cmrA* is required for PleD activation, we would expect combining the Δc*mrA* and Δ*dgcB* mutations to have a synergistic effect that leads to a complete loss of adhesion. Alternatively, if *cmrA* is required for DgcB activation, a Δ*cmrA* Δ*pleD* double mutant should be completely non-adhesive. We first determined if *cmrA* is required for activating PleD or DgcB in the Δ*cheYII* background. Deleting *cmrA* suppressed adhesion more strongly than deletion of *pleD* or *dgcB* in the Δ*cheYII* background. Adhesion in the Δ*cheYII* Δ*pleD* Δ*cmrA* triple mutant was indistinguishable from the Δ*cheYII* Δ*cmrA* double mutant. The Δ*cheYII* Δ*dgcB* Δ*cmrA* triple mutant was completely non-adhesive (Fig. 3A) and indistinguishable from the Δ*cheYII* Δ*dgcB* Δ*pleD* mutant. The synergistic effect of mutating *cmrA* and *dgcB* indicates that *cmrA* is required for activation of PleD in the Δ*cheYII* mutant background.

**Figure 3.**
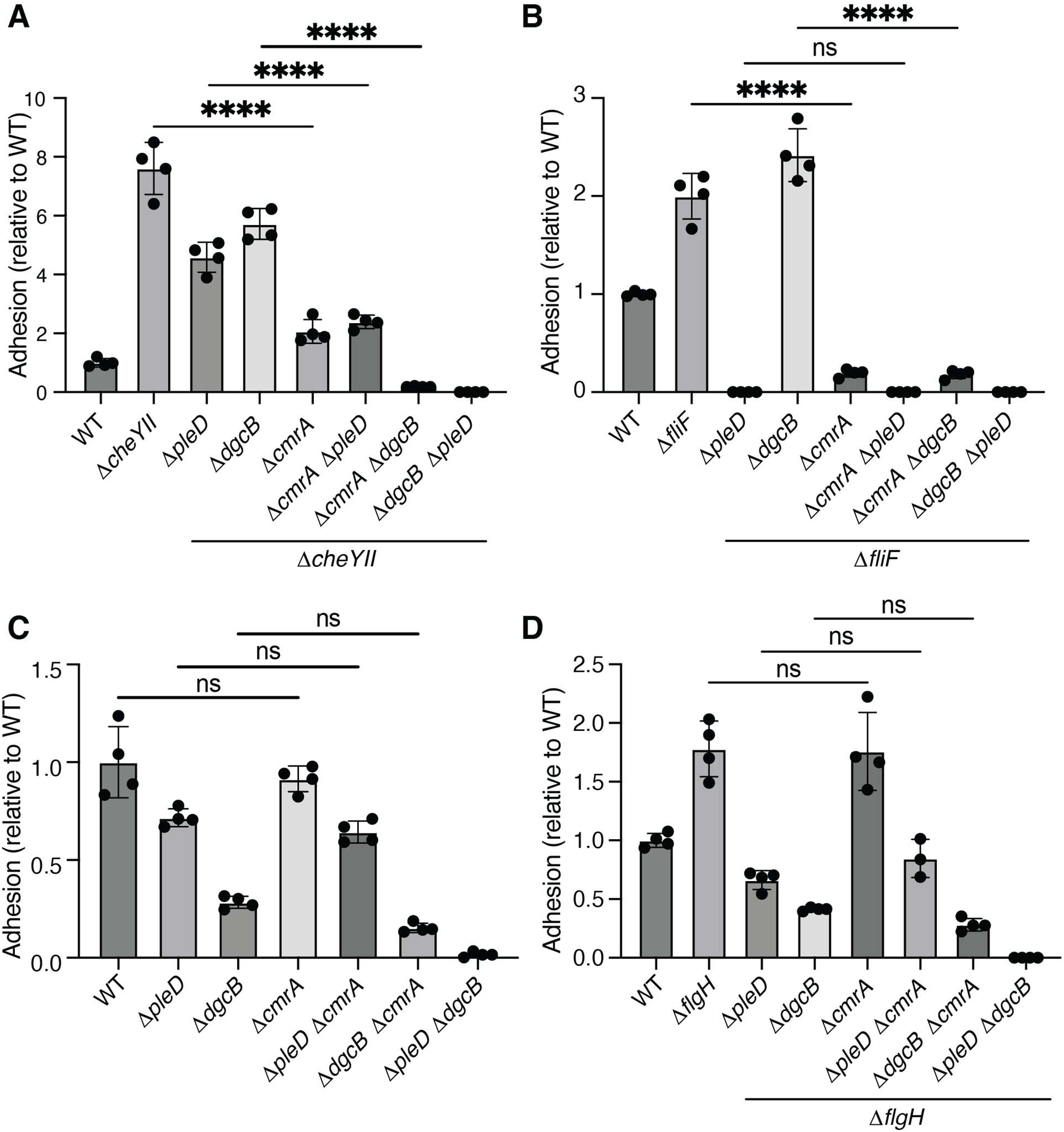
*cmrA* effects *pleD* signaling in a context-dependent manner. Representative crystal violet (CV) staining to measure adhesion of Δ*cmrA* in the (a) Δ*cheYII* background, (b) Δ*fliF* background, (c) WT background, and (d) Δ*flgH* background. Three biological replicates were performed for each CV stain, and each replicate contained four technical replicates. Statistical significance was determined by one-way ANOVA test and Tukey’s multiple-comparison test. Four asterisks indicate a P-value < 0.0001.

We next tested the role of *cmrA* in the Δ*fliF* background where *dgcB* is dispensable and adhesion is completely dependent on *pleD*. If *cmrA* plays a role in *pleD-*driven hyperadhesion, we would expect the Δ*fliF* Δ*cmrA* double mutant to exhibit weak adhesion compared to Δ*fliF* alone. Indeed, the Δ*fliF* Δ*cmrA* strain exhibited extremely low levels of adhesion, supporting the conclusion that *cmrA* is involved in *pleD* regulation (Fig. 3B). The result that *cmrA* plays a key role in Δ*fliF* hyperadhesion is consistent with our findings in the Δ*cheYII* background in Fig. 3A.

The importance of *cmrA* in PleD regulation is context specific. Deleting *cmrA* had no effect on adhesion in the wild-type, Δ*pleD* or Δ*dgcB* backgrounds (Fig. 3C). Likewise, introducing *cmrA* deletions into the Δ*flgH,* Δ*flgH* Δ*pleD*, or Δ*flgH* Δ*dgcB* mutants had no effect on adhesion (Fig. 3D). The lack of an additive effect between the Δ*cmrA* and Δ*dgcB* mutations indicates that *cmrA* is not involved in activating PleD in the wild-type or Δ*flgH* backgrounds. Taken together, our epistasis experiments support the model that *cmrA* is required for *pleD-*dependent adhesion activation in the Δ*fliF* and Δ*cheYII* backgrounds but not in the wild-type or Δ*flgH* backgrounds.

### Context dependent localization of CmrA-Venus to the flagellar pole

The distinct effects of disrupting *cmrA* in different flagellar mutant backgrounds suggested that *cmrA* responds to specific assembly states of the flagellar motor. We hypothesized that CmrA may be recruited to the flagellum in response to certain disruptions to flagellar function. We tagged CmrA by fusing *venus* to the 3’ end of *cmrA* at the native chromosomal locus to generate a C-terminal fusion. The resulting CmrA-Venus protein was functional, as the Δ*cheYII* mutant remained hyperadhesive when *cmrA* was replaced by *cmrA-venus* (Fig. S1). In wild-type cells, we observed a predominantly diffuse signal in the cytoplasm with occasional CmrA-Venus foci at a single pole. We analyzed 1077 WT cells and determined that only 10% of cells exhibited polar foci. Additionally, 24% of WT cells contained non-polar foci. We performed timelapse microscopy and determined that CmrA-Venus foci form at the flagellar pole in swarmer cells, but the foci can remain into the swarmer to stalked cell transition phase (Fig. S2).

Because *cmrA* has no obvious phenotype in the WT background (Fig. 2D, Fig. S3) we also measured CmrA-Venus localization in the Δ*cheYII* background in which *cmrA* is involved in promoting adhesion. We observed a sharp increase in the number of cells with polar CmrA localization and a corresponding decrease in cells with diffuse or non-polar signal (Fig. 4A). Of the 730 Δ*cheYII* cells analyzed, 36% exhibited polar localization, and only 6% exhibited non-polar localization. Compared to WT, this is a 3-fold increase of polar localization and a 4-fold decrease in non-polar localization (Fig. 4C). This analysis confirms that CmrA-Venus localizes to the pole more strongly in Δ*cheYII* than in WT.

**Figure 4.**
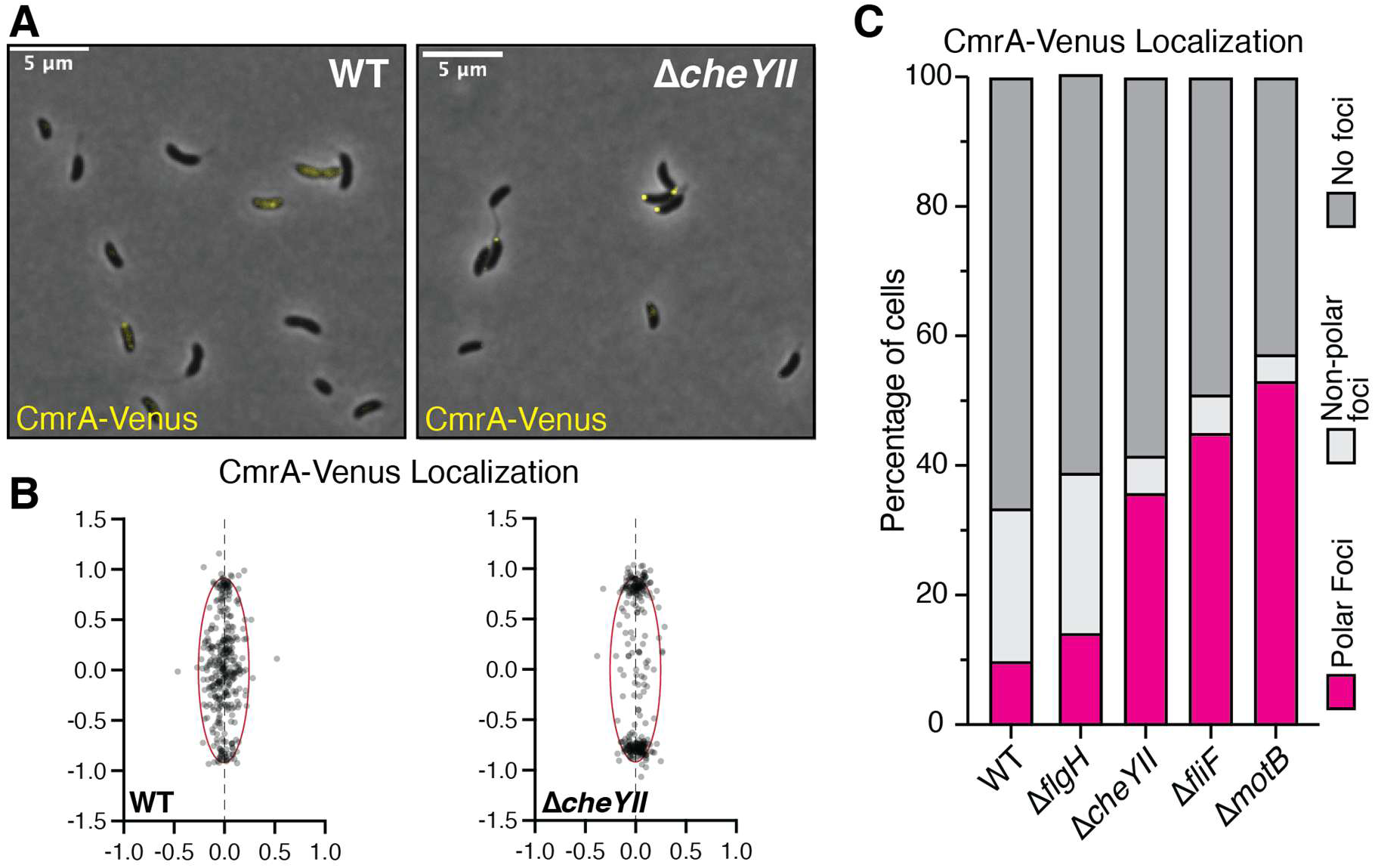
CmrA-Venus localizes to the pole in specific flagellar assembly mutant backgrounds. (A) Representative micrographs of CmrA-Venus in WT and Δ*cheYII* backgrounds. (B) Compilation of all foci on a representative cell to visualize general localization of foci. (C) Stacked bar graph showing the percentage of cells with polar CmrA-Venus foci in each genetic background. WT n=1077, Δ*flgH* n=1066, Δ*cheYII* n=730, Δ*fliF* n=963, Δ*motB* n=942.

Increased polar localization of CmrA-Venus in the Δ*cheYII* mutant suggested that recruitment to the pole may occur specifically in genetic backgrounds where *cmrA* contributes adhesion. To test if flagellar assembly influences CmrA localization, we analyzed CmrA-Venus in the Δ*flgH* and Δ*fliF* backgrounds. We found that 45% of Δ*fliF* cells exhibited a polar focus compared to only 14% of Δ*flgH* cells. Because CmrA localized in the flagellum-null strain (Δ*fliF*) but remained largely diffuse in the presence of a partially assembled structure containing a C-ring (Δ*flgH*), we reasoned that CmrA might be recruited in response to altered rotation of the C-ring by the MotAB stators. Indeed, 53% of cells exhibited a polar focus in a Δ*motB* mutant that contains a fully assembled but rotationally paralyzed flagellum. These results indicate that specific rotational states of the flagellar motor promote CmrA recruitment to the pole.

### Altered directional switching promotes polar recruitment of CmrA

The finding that CmrA localization was enhanced in the Δ*cheYII,* Δ*fliF* and Δ*motB* mutant backgrounds suggested that altering the directional rotation of the C-ring may activate *cmrA* signaling. To test this model, we first examined CmrA localization in double mutants combining the Δ*flgH* deletion with either the Δ*cheYII* or Δ*motB* mutations. We predicted that disrupting the rotational switching of the C-ring would drive CmrA to the pole even in the presence of a partially assembled HBB. Deleting *cheYII* or *motB* enhanced the polar localization of CmrA in the Δ*flgH* background (Fig. 5A). Compared to the Δ*flgH* background where only 9% of cells contained polar foci, 27% of Δ*flgH* Δ*cheYII* and 34% of Δ*flgH* Δ*motB* cells contained polar foci. These data provide further support for the model that the rotational status of the motor rather than the assembly alone controls recruitment of CmrA to the cell pole.

**Figure 5.**
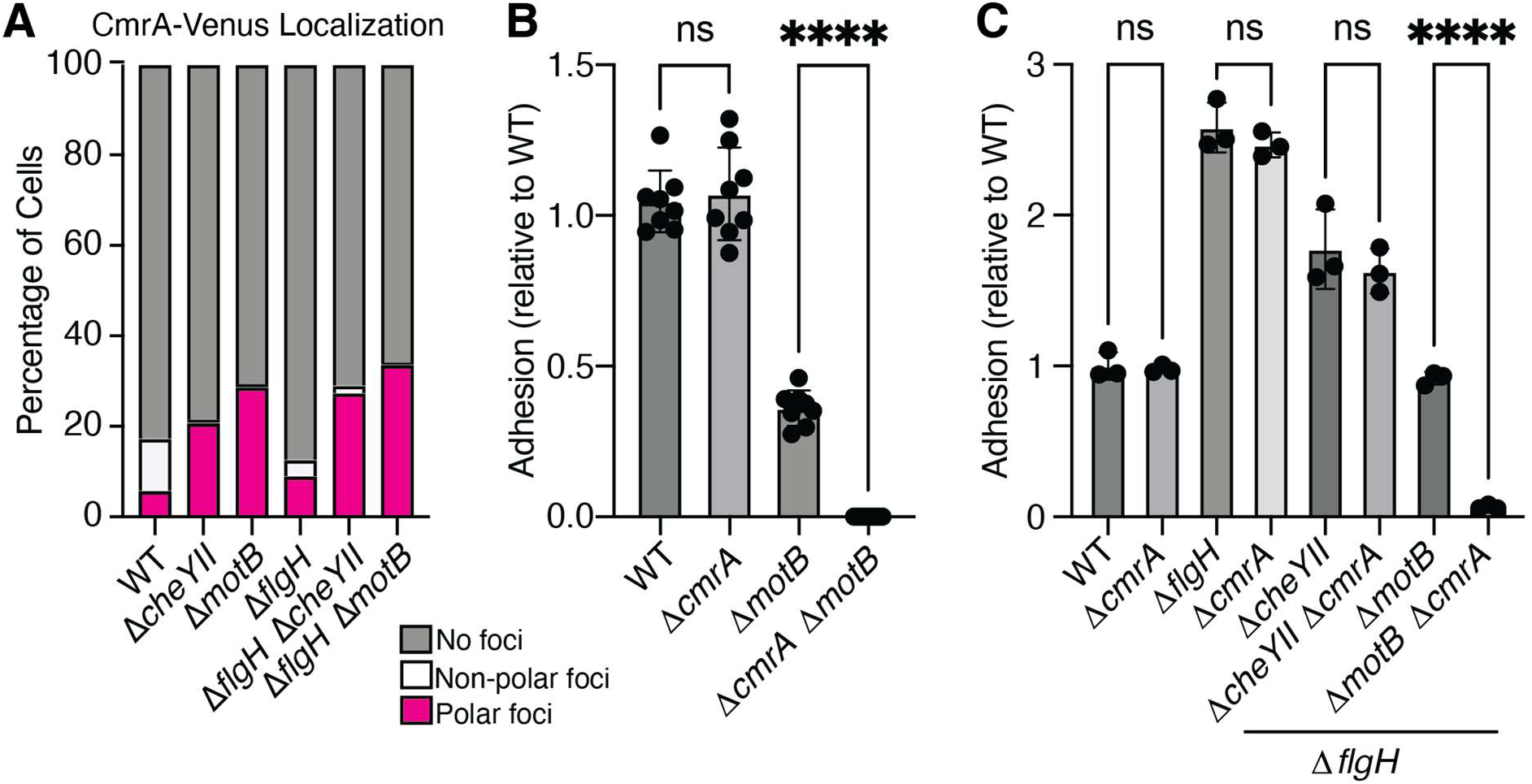
Disruption of the stators makes adhesion dependent on *cmrA.* Stacked bar graph representing the percentage of CmrA-Venus foci in each genetic background. WT n=808, Δ*cheYII* n=807, Δ*motB* n=624, Δ*flgH* n=1221, Δ*flgH* Δ*cheYII* n=830, Δ*flgH* Δ*motB* n=764. (B and C) Representative crystal violet (CV) staining to measure adhesion. Statistical significance was determined by one-way ANOVA test and Tukey’s multiple-comparison test. Four asterisks indicate a P-value < 0.0001.

We next tested whether enhanced CmrA polar localization drives the requirement of *cmrA* for adhesion using CV staining assays. Given the high polar localization of CmrA-Venus in the Δ*motB* mutant, we predicted that *cmrA* would contribute to adhesion in this background. Indeed, a Δ*cmrA* Δ*motB* double mutant was completely non-adhesive (Figure 5), showing that *cmrA* signaling is active in the Δ*motB* background. We then tested if blocking directional switching would render the Δ*flgH* mutant dependent on *cmrA* for adhesion. Adhesion in the Δ*flgH* Δ*motB* background was entirely dependent on *cmrA*. In contrast, deleting *cmrA* in the Δ*flgH* Δ*cheYII* background had no effect as a Δ*flgH* Δ*cheYII* Δ*cmrA* triple mutant was indistinguishable from the double mutant. We conclude that altering the directional rotation of the C-ring drives CmrA to the cell pole independent of complete flagellum assembly but that the enhanced localization of CmrA-Venus is not sufficient for adhesion regulation.

### CmrA is a degenerate GGDEF/EAL domain protein

*cmrA* was annotated as a hypothetical protein with no functional prediction. However, a structural prediction (30) indicated that CmrA is a tandem GGDEF/EAL domain protein (Figure 6A). The predicted active sites of both domains contained many degenerate residues, suggesting that CmrA does not affect c-di-GMP metabolism directly. Instead, we investigated whether residues known to contribute to nucleotide binding were required for CmrA activity. We mutated 10 residues to alanine across both the predicted GGDEF and EAL domains. We fused *cmrA* mutant alleles to the native *cmrA* promoter, inserted them at the xylose locus and tested their functionality in the Δ*cheYII* Δ*cmrA* background. If a mutant allele restores Δ*cheYII* hyperadhesion in the Δ*cheYII* Δ*cmrA* background, then that residue is not necessary for CmrA function. Alternatively, if the mutant allele is unable to restore hyperadhesion, then that residue is necessary for functionality. We found that many of the charged residues in both the predicted GGDEF domain and EAL domain were important for CmrA function (Figure 6C).

**Figure 6.**
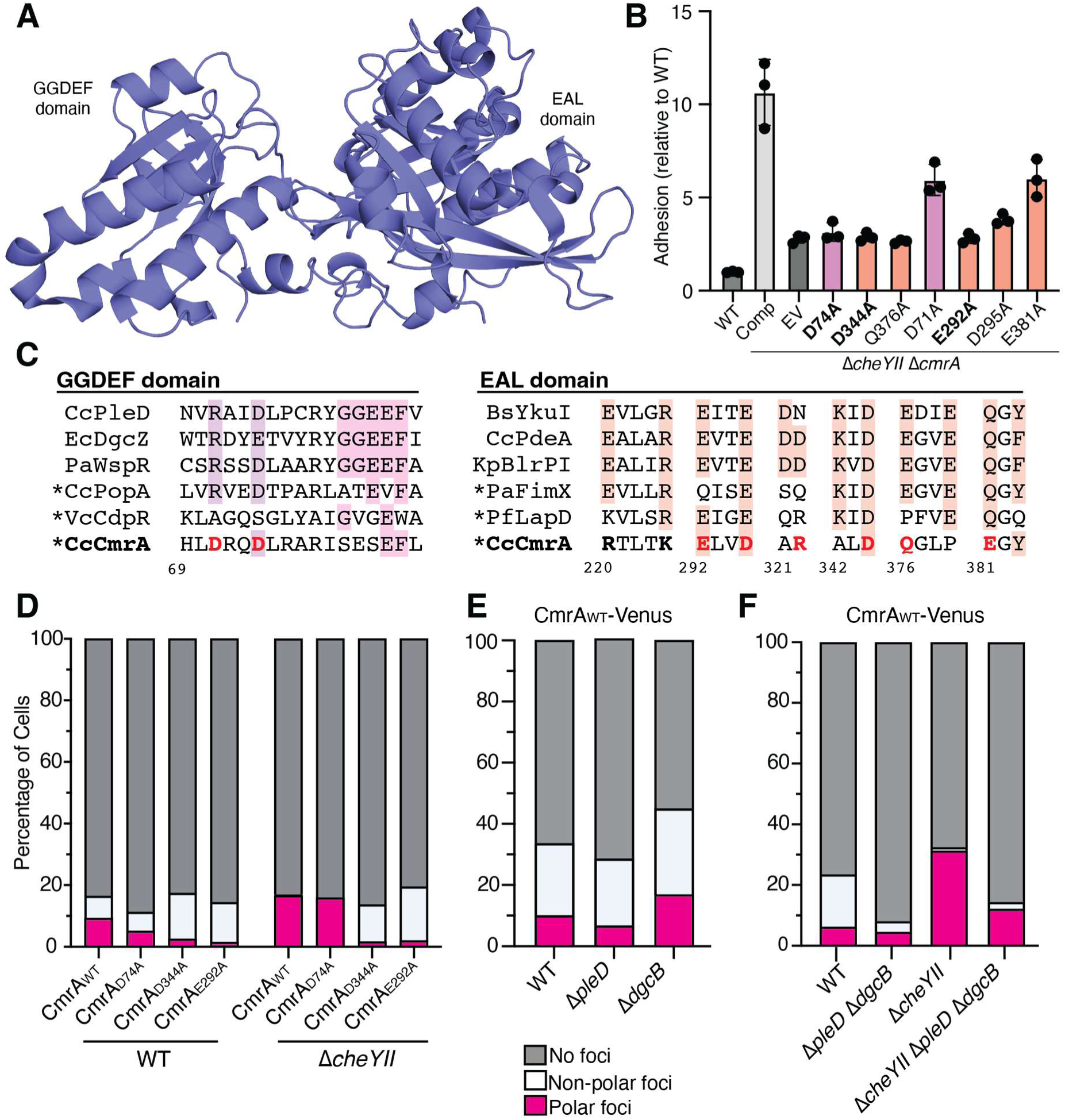
CmrA is a predicted c-di-GMP binding protein. (A) Alpha fold model of CmrA showing a predicted tandem GGDEF/EAL domain structure. (B) CV Staining comparing complementation of *cmrA* in a Δ*cmrA* Δ*cheYII* background with CmrA mutant alleles. (C) Alignment of CmrA with other GGDEF and EAL domain proteins. Important residues are highlighted. Bolded residues were mutated in CmrA. Red bolded residues were required for full hyperadhesion in Δ*cheYII* (D) Quantification of percentage of cells containing polar foci of CmrA-Venus point mutants in a WT or Δ*cheYII* background. WT n=810, CmrA_D74A_ n=848, CmrA_D344A_ n=887, CmrA_E292A_ n=1399, Δ*cheYII* n=1450, Δ*cheYII* CmrA_D74A_ n=790, Δ*cheYII* CmrA_D344A_ n=917, Δ*cheYII* CmrA_E292A_ n=1025 (E&F) CmrA-Venus localization in Δ*pleD* and Δ*dgcB* backgrounds.

We first examined charged residues in the GGDEF domain that we predict comprise the I-site of the protein. The I-site serves as a non-competitive, allosteric binding site for dimerized c-di-GMP and is comprised of an RXXD sequence upstream of the GGDEF motif (31, 32). We mutated residues D71 and D74 of the modified DXXD sequence at the I-site positions in the CmrA GGDEF domain. Neither allele restored CmrA activity in the Δ*cheYII* Δ*cmrA* background, but CmrA_D74A_ was less functional than CmrA_D71A_ (Figure 6B). We also examined EAL domains residues that contribute to substrate binding and ion coordination (33). The degenerate ExLxR sequence for which the domain is named was dispensable for CmrA function (Fig. S4). However, alleles with substrate binding (D344, Q376) and Mg2+ ion coordination (E292) residues mutated were unable to restore hyperadhesion in the Δ*cheYII* Δ*cmrA* background.

We hypothesized that mutant CmrA proteins unable to support Δ*cheYII* hyperadhesion may be defective in localizing to the cell pole. We examined the localization of mutant CmrA-Venus fusions expressed from the native *cmrA* locus. The CmrA_E292A_ and CmrA_D344A_ mutants were unable to localize to the pole in the WT background or in the Δ*cheYII* background (Fig. 6D). Immunoblotting confirmed that the localization patterns were not caused by reduced stability of CmrA (Fig S5), and CV staining confirmed that neither CmrA_E292A_-Venus nor CmrA_D344A_-Venus supported Δ*cheYII* hyperadhesion (Figure S5). We performed the same analysis for CmrA_D74A_ and found that it retained its ability to localize to the cell pole (Fig. 6D), but the Venus-tagged allele also supported hyperadhesion in the Δ*cheYII* background (Fig S4). Because the adhesion phenotype of the *cmrA_D74A_*-venus allele differs from its untagged counterpart, we cannot definitively conclude whether residue D74 affects polar localization. Nevertheless, these results indicate that conserved residues within the GGDEF I-site and EAL domain are required for CmrA function.

Because structural modeling and mutagenesis indicated that nucleotide-binding residues are important for CmrA function, we considered whether c-di-GMP regulates CmrA localization. To test this, we measured localization of CmrA-Venus in the Δ*pleD* and Δ*dgcB* backgrounds (Figure 6E). In the Δ*pleD* background, 7% of cells contained a polar CmrA-Venus focus, while 16% of Δ*dgcB* cells contained a polar focus. These localization patterns were similar to wild type and indicate that neither DGC is specifically required for CmrA recruitment to the cell pole. We then measured CmrA localization in the double Δ*pleD* Δ*dgcB* deletion and in a Δ*pleD* Δ*dgcB* Δ*cheYII* triple deletion. Despite the fact that cells were elongated and stalk development was clearly disrupted in Δ*pleD* Δ*dgcB* double mutant, 5% of cells retained a polar CmrA-Venus focus and 3% exhibited non-polar localization. In the Δ*pleD* Δ*dgcB* Δ*cheYII* triple deletion mutant, 12% of cells exhibited a polar localization pattern and 2% exhibited non-polar localization. These data suggest that CmrA-Venus can still respond to perturbations in the motor and localize to the cell pole in the absence of c-di-GMP signaling.

## Discussion

The flagellum is a key surface sensing apparatus in many bacteria. Current models propose that when cells encounter a surface the resulting obstruction of flagellar rotation stimulates c-di-GMP production and biofilm formation (1). The increased load from obstruction of flagellar rotation is thought to activate downstream signaling pathways that promote adhesion (34, 35). However, we and others have shown that the motor also regulates attachment by incorporating signals from directional switching that occurs during chemotaxis (16, 18, 36, 37). In *C. crescentus*, surface sensing pathways can be stimulated by disrupting the assembly (Δ*fliF* and Δ*flgH*) or directional switching activity (Δ*cheYII*) of the flagellum, resulting in differential activation of the diguanylate cycases DgcB and PleD. DgcB activation is thought to require functional MotAB stators, while PleD activation is thought to be stator-independent (9). Our previous study challenged this model by showing that MotAB stators are linked to PleD activation specifically in chemotaxis mutants. In this paper, we used a genetic screen to identify a previously uncharacterized gene (CCNA_02061, renamed *cmrA*) that is required for hyperadhesion in a chemotaxis (Δ*cheYII*) mutant background. Our results indicate that *cmrA* is required for PleD activation when flagellar motor rotation is disrupted.

The canonical model for PleD regulation involves the opposing effects of two histidine kinases. PleC acts as a phosphatase to maintain low PleD activity at the swarmer cell pole, and DivJ acts as a kinase to activate PleD at the incipient stalked cell pole (27). We leveraged the synergistic loss of adhesion observed when PleD and DgcB are inactivated simultaneously to demonstrate that *cmrA* represents a novel, context-dependent factor in PleD signaling. Combining the Δ*cmrA* and Δ*dgcB* mutations has a synergistic effect in the Δ*cheYII* background that leads to a loss of adhesion (Fig 3). This result demonstrates that *cmrA* is required for *pleD* function in chemotaxis mutants. However, the association between *cmrA* and *pleD* is context dependent. No synergistic interaction between the *cmrA* and *dgcB* mutations was observed in the wild-type or Δ*flgH* backgrounds, even though *pleD* promotes adhesion under these conditions. Thus, PleD activates adhesion in a *cmrA-*dependent or -independent manner depending on the functional state of the flagellum.

Although CmrA clearly contributes to PleD function when chemotaxis is disrupted, our epistasis data provide conflicting indications as to whether CmrA acts upstream to activate PleD or as a downstream effector. From a strictly genetic perspective, mutating Δ*cmrA* suppresses adhesion more strongly than the Δ*pleD* mutation in the Δ*cheYII* background, and the Δ*cheYII* Δ*cmrA* Δ*pleD* triple mutant phenocopies the more strongly suppressed hyperadhesion of the Δ*cheYII* Δ*cmrA* mutant. This epistatic relationship places *cmrA* downstream of *pleD* in the signaling cascade. However, in the Δ*fliF* background, the Δ*cmrA* mutation suppresses adhesion less than the Δ*pleD* mutation, placing *cmrA* upstream of *pleD*. It is possible that the difficulty in determining if CmrA is an activator of PleD or a PleD-dependent effector is indicative of the relationship between the two factors. c-di-GMP effectors and DGCs such as PleD are often regulated by feedback loops, and we speculate that PleD-dependent activation of CmrA causes further enhancement of PleD signaling to amplify surface responses (31, 32). Feedback loops are common features of c-di-GMP signaling in other organisms as well. For example, in *Pseudomonas aeruginosa,* the flagellar stators and the DGC SadC form a positive feedback loop resulting in enhancement of c-di-GMP levels and motor disassembly (38). The *P. aeruginosa* c-di-GMP effector protein LapD (39) binds to c-di-GMP creating a conformational change that allows the DGC GcbC to bind to LapD. This interaction is mediated by the binding of citrate and enhances GcbC activity (40–42). Though we cannot say with certainty that CmrA acts downstream of PleD, a connection between the two genes is clear from our epistasis experiments.

CmrA has a predicted tandem GGDEF/EAL domain structure but is unlikely to be directly involved in c-di-GMP metabolism due to its degenerate GGDEF and EAL motifs. While we have so far been unable to purify the protein for *in vitro* studies, we showed that mutating residues that affect c-di-GMP binding in other GGDEF/EAL domain proteins can disrupt the protein’s regulatory function. A *cmrA* allele with a mutation of the I-site (D74A) upstream of the predicted GGDEF sequence is unable to support hyperadhesion. Interestingly, the CmrA_D74A_-Venus allele restores hyperadhesion and localizes to the cell pole in the Δ*cheYII* background. These data indicate that mutation of the I-site may not completely abolish CmrA function. It is possible the fusion of the Venus tag to the C-terminus protects CmrA from degradation. The C-terminus of CmrA consists of three alanine residues, which is a well-documented degradation tag for the ClpXP protease (43). This might explain why the untagged allele is not able to restore Δ*cheYII* hyperadhesion, but the tagged allele appears fully functional. Mutating residues that contribute to c-di-GMP binding in other EAL-domain proteins (E292 and D344) disrupts both the role of *cmrA* in Δ*cheYII* hyperadhesion and the polar localization of CmrA-Venus, even in conditions that favor polar localization of the CmrA_WT_-Venus allele. These results demonstrate that potential ligand-binding residues within both the GGDEF and EAL domains are essential for CmrA functionality. These results suggest that CmrA acts as a c-di-GMP effector protein that actuates PleD-signaling.

CmrA-Venus is recruited to the cell pole in a specific subset of flagellar mutant backgrounds. Its localization is enhanced when directional switching of the flagellar motor is altered through loss of the C-ring, CheYII, or the stators but not in the wild-type or Δ*flgH* backgrounds. Though its localization depends on the assembly state of the flagellum, CmrA is unlikely to be recruited directly by the flagellum because it retains enhanced polar localization in the Δ*fliF* mutant in which the earliest stage assembly is blocked. Furthermore, CmrA-Venus localization is highest during the transition between the swarmer and stalked cell stages when flagellar disassembly is occurring. It is possible that CmrA interacts with another protein that localizes to the pole when the motor is interrupted. Alternatively, directional switching may influence liquid-liquid phase separation at the cell pole. In this model, disrupting the rotation of the motor would promote condensate formation at the pole that traps CmrA. There are many examples of phase separation at the poles driving cellular differentiation in *C. crescentus* (44–46). These two possibilities are not mutually exclusive, and further studies are needed to determine the mechanism of CmrA’s recruitment to the cell pole.

Collectively, our findings emphasize how the flagellar motor functions as a multi-state signaling hub rather than a simple transducer of mechanical load. Structural variations such as the complete absence of the flagellum (Δ*fliF*), partial complex assembly (Δ*flgH*) and locked rotational direction (Δ*cheYII*) each activate discrete c-di-GMP signaling pathways. Disentangling the dense, overlapping topology of these behavioral states presents a significant genetic challenge, as critical regulatory factors like CmrA can remain entirely hidden under standard screening conditions. The genome-wide adhesion profiling approach has proven ideal for resolving this network architecture. By contrasting the fitness profiles of mutant libraries across a panel of flagellar signaling backgrounds, we were able to reduce genetic redundancy and isolate a context-specific pathway. Utilizing differential high-throughput screens as a genetic prism to separate distinct operational states of a single multi-protein machine represents a powerful framework for dissecting multi-layered, state-dependent signaling networks across diverse bacterial species.

## Materials and Methods

### Bacterial growth and genetic manipulations

*Escherichia coli* strains were grown at 37°C in LB medium supplemented with 1.5% (w/v) agar, 50μg/mL kanamycin, 300μM diaminopimelic acid (DAP) and agitation at 200rpm when necessary. Unless otherwise indicated, *C. crescentus* was grown in peptone yeast extract (PYE) broth. Cultures were grown at 30°C with 1.5% (w/v) agar, 3% (w/v) sucrose, 25μg/mL kanamycin (solid medium), 5μg/mL kanamycin (liquid medium) and shaking at 200 rpm when necessary. Purified plasmids were mobilized into *C. crescentus* using electroporation. All *C. crescentus* experiments were performed in the CB15 strain background.

Strains and plasmids used in this study are listed in Tables S2 and S3, respectively. Plasmids were either synthesized commercially through the Azenta Genewiz ValueGene service or developed by PCR amplification and subsequent Gibson assembly into restriction digested plasmids. All plasmids were verified by complete sequence assembly service (Plasmidsaurus). Plasmids for generating gene deletions were constructed by fusing ∼500bp upstream and ∼500bp downstream of the gene of interest with the first 4 codons (12bp) and the last 4 codons (12 bp) including the stop codon at the junction between the fragments. The plasmid for fusing *venus* to *cmrA* was constructed by fusing 500bp upstream of the stop codon, the Venus open reading frame and 500bp downstream stop codon. Genetic complementation of the Δ*cmrA* mutation was performed by integrating the *cmrA* open reading frame under the control of the native *cmrA* promoter at the *xyl* locus using pXGFPC-2. The *PcmrA-cmrA* cassette was inserted in reverse orientation to avoid interference from the *xyl* promoter. Mutant *cmrA* alleles were generated using the Agilent QuickChange protocol with a subsequent Gibson assembly performed on the Dpn1-treated product to increase yields of positive clones.

### Crystal violet staining

Strains grown overnight in 2mL of PYE and backdiluted to an OD_660_ of 0.5 in PYE. Individual wells of a 48-well plate were then filled with 450uL of M2X media (1X M2 387 salts, 1% hunter base, 0.5 M CaCl2, 1M MgSO4, and 0.15% xylose), and each well was inoculated with 1.5uL of backdiluted culture. Each plate contained 3-4 technical replicates of each strain. Plates were incubated at 30°C for 17 hours shaking in an orbital shaker at 150 rpm. Cultures were discarded, and plates were then thoroughly washed with tap water. Each well was incubated with 500uL of 0.01% crystal violet for 10 minutes shaking at RT at approximately 150 rpm. The crystal violet stain was discarded, and the plate was once again washed thoroughly with tap water. 500uL of ethanol was then added to each well and plates were incubated again for approximately 10 minutes. Quantification of the CV dye was performed by measuring the absorbance at 575 nm using a 396 BioTek Synergy H1microplate reader. The absorbance value from wells containing only M2X was then subtracted from the sample absorbance values as background. Each plate contained WT *Caulobacter crescentus* as an internal control. Absorbance values of each well were normalized to the mean value of the WT samples to calculate the relative levels of adhesion.

### RB-TnSeq

A randomly barcoded transposon library was constructed in the Δ*cheYII* background as described previously (47) based on the method of (28). Briefly, mid-log phase cultures of the APA_752 barcoded transposon pool and *C. crescentus* Δ*cheYII* mutant were collected, with PYE medium containing DAP, mixed and spotted on a PYE agar plate containing DAP overnight. Cells were collected, homogenized in PYE medium and spread on PYE plates containing kanamycin. After 72 hours of growth, colonies were scraped from plates, pooled, outgrown, aliquoted and frozen in 15% (v/v) glycerol.

Genomic DNA was isolated and sheared to generate ∼300bp fragments. Transposon junctions were amplified using the primers TS_pHimar_f (5’-ACACTCTTTCCCTACACGACGCTCTTCCGATCTCGCCCTGCAGGGATGTCCACGAG-3’) and TS_r (5’-GTGACTGGAGTTCAGACGTGTGCTCTTCCGATCT-3’). The amplification product was purified and subjected to dual indexing PCR. 150bp paired end reads were collected on an Illumina NovaSeq X Plus sequencer. Reads were mapped and used to generate a list of barcoded insertions using the MapTnSeq.pl and DesignRandomPool.pl scripts (28). Sequencing data from the RB-TnSeq experiment has been deposited in the NCBI Sequence Read Archive as BioProject PRJNA1467683.

### Adhesion profiling with cheesecloth

Adhesion profiling was performed as described previously (8). Aliquots of the barcoded Δ*cheYII* transposon library were serially passaged in 12-well microtiter plates containing 500μL M2X medium and a patch of cheesecloth for five passages. An aliquot of the spent medium from each passage was centrifuged and the cell pellets were used as templates for barcode amplification using the primers TS_U1 (5’-ACACTCTTTCCCTACACGACGCTCTTCCGATCTGATGTCCACGAGGTCTCT-3’) and TS_U2 (5’-GTGACTGGAGTTCAGACGTGTGCTCTTCCGATCTGTCGACCTGCAGCGTACG-3’). Amplification products were purified and subjected to dual indexing PCR. 150bp paired end reads were collected on an Illumina NovaSeq X Plus sequencer. Barcodes were scored using the MultiCodes.pl and CombineBarSeq.pl scripts, and fitness values were determined using the BarSeqR.pl script (https://bitbucket.org/berkeleylab/feba/src/master/). Sequencing data from the BarSeq experiment has been deposited in the NCBI Sequence Read Archive as BioProject PRJNA1467683.

### Fluorescence microscopy

Strains were grown overnight in 2mL of PYE. Cultures were backdiluted 1:10 in M2X medium and grown to an OD_660_ of 0.2-0.4. CmrA-Venus fusion strains were backdiluted in PYE. Slides were set up with a thin 3% (w/v) agarose pad (∼135uL), made with low melting point agarose. 2uL of culture were spotted on the agarose pad and finished with a coverslip. Slides were imaged on a Nikon Eclipse Ti series inverted microscope with a 100x oil immersion objective lens. Fluorescence images were collected using a Prior 424 Lumen 200 metal halide light source and a YFP-specific filter set (Chroma). Images were taken with 25-50 ms phase exposure and 2s fluorescence exposure. Images were processed using ImageJ, and MicrobeJ (48) was used to quantify the number of cells with polar foci. Strains from each panel were all imaged during the same microscopy session and analyzed using identical image processing parameters.

### Timelapse microscopy

Strains were grown overnight in PYE medium and backdiluted 10-fold in PYE. Cells were spotted on a 1.5% (w/v) agarose pad made with low melting point agarose. Pads were sealed and placed at 30°C to incubate for 20 minutes before imaging. Slides were imaged for 3 hours with images taken every 30 minutes with 50ms phase exposure and 500ms YFP exposure on a Nikon Eclipse Ti series inverted microscope with a 100x oil immersion objective lens.

### Immunoblotting

Cultures were grown in 2mL of PYE overnight and backdiluted 10-fold in M2X medium. Cells were harvested at an OD_660_ of 0.2-0.4. A 1.0 OD_660_ equivalent of cells was centrifuged for 1 minute at 3500 x g followed by an additional 2 minutes at 17000 x g. Supernatant was decanted and cell pellets were resuspended in 50mL of TBS. 1uL of turbonuclease (Sigma) was added to reduce viscosity in samples. Samples were then incubated at 30°C for 10 minutes, and 25uL of 4x Laemelli buffer was added to each sample before incubating at 100°C for 10 minutes. Samples were cooled at room temperature and stored −20°C. Samples were heated to 65°C and then loaded into a 12% acrylamide mini-gel, resolved and transferred to a PVDF membrane via semi-dry transfer. Blocking was performed with 5% (w/v) milk powder in TBST at room temp for 2 hours. Membranes were incubated overnight at 4°C in a solution of 5% milk powder containing a 1:10000 dilution of anti-GFP serum from rabbit. Washes were then performed in TBST and the membrane was incubated in 5% milk powder containing 1:10000 dilution of goat anti-rabbit HRP for 90 minutes at room temperature. Another round of washes in 1x TBST was performed and the membrane was briefly incubated in 1x TBS before soaking the membrane in Western Lightning Plus substrate (PerkinElmer). Membranes were imaged on Invitrogen iBright FL1500 Imaging System.

## Data availability

Illumina sequencing data was uploaded to the Sequence Read Archive under BioProject PRJNA1467683.

## Acknowledgements

We thank Chandler Hellenbrand for critical reading of the manuscript. This work was funded by National Institutes of Health (NIH) grant 1R35GM150652 to D.M.H.

## References

1. Berne C, Ellison CK, Ducret A, Brun YV. 2018. Bacterial adhesion at the single-cell level. Nat Rev Microbiol 16:616–627.

2. Muhammad MH, Idris AL, Fan X, Guo Y, Yu Y, Jin X, Qiu J, Guan X, Huang T. 2020. Beyond Risk: Bacterial Biofilms and Their Regulating Approaches. Front Microbiol 11:928.

3. Li G, Brown PJB, Tang JX, Xu J, Quardokus EM, Fuqua C, Brun YV. 2012. Surface contact stimulates the just-in-time deployment of bacterial adhesins. Molecular Microbiology 83:41–51.

4. Belas R. 2014. Biofilms, flagella, and mechanosensing of surfaces by bacteria. Trends Microbiol 22:517–527.

5. Laventie B-J, Jenal U. 2020. Surface Sensing and Adaptation in Bacteria. Annu Rev Microbiol 74:735–760.

6. Ellison CK, Kan J, Dillard RS, Kysela DT, Ducret A, Berne C, Hampton CM, Ke Z, Wright ER, Biais N, Dalia AB, Brun YV. 2017. Obstruction of pilus retraction stimulates bacterial surface sensing.

7. Hug I, Deshpande S, Sprecher KS, Pfohl T, Jenal U. 2017. Second messenger–mediated tactile response by a bacterial rotary motor. Science 358:531–534.

8. Hershey DM, Fiebig A, Crosson S. 2019. A Genome-Wide Analysis of Adhesion in Caulobacter crescentus Identifies New Regulatory and Biosynthetic Components for Holdfast Assembly. mBio 10:e02273–18.

9. Hershey DM, Fiebig A, Crosson S. 2021. Flagellar Perturbations Activate Adhesion through Two Distinct Pathways in Caulobacter crescentus. mBio 12:e03266–20.

10. Wu J, Newton A. 1997. Regulation of the Caulobacter flagellar gene hierarchy; not just for motility. Molecular Microbiology 24:233–239.

11. Chevance FFV, Hughes KT. 2008. Coordinating assembly of a bacterial macromolecular machine. Nat Rev Microbiol 6:455–465.

12. Einenkel R, Halte M, Sawant SA, Erhardt M, Wadhwa N, Popp PF. 2025. Building the bacterial flagellum: coordinating regulation, dynamic assembly, and function. Microbiology and Molecular Biology Reviews 89:e00092–22.

13. Santiveri M, Roa-Eguiara A, Kühne C, Wadhwa N, Hu H, Berg HC, Erhardt M, Taylor NMI. 2020. Structure and Function of Stator Units of the Bacterial Flagellar Motor. Cell 183:244–257.e16.

14. Deme JC, Johnson S, Vickery O, Aron A, Monkhouse H, Griffiths T, James RH, Berks BC, Coulton JW, Stansfeld PJ, Lea SM. 2020. Structures of the stator complex that drives rotation of the bacterial flagellum. Nat Microbiol 5:1553–1564.

15. Bi S, Sourjik V. 2018. Stimulus sensing and signal processing in bacterial chemotaxis. Current Opinion in Microbiology 45:22–29.

16. Salemi RI, Cruz AK, Hershey DM. 2024. A flagellar accessory protein links chemotaxis to surface sensing. J Bacteriol 206:e0040424.

17. Zappa S, Berne C, Morton Iii RI, Whitfield GB, De Stercke J, Brun YV. 2024. The HmrABCX pathway regulates the transition between motile and sessile lifestyles in *Caulobacter crescentus* by a mechanism independent of *hfiA* transcription. mBio e01002–24.

18. Laganenka L, Lee J-W, Malfertheiner L, Dieterich CL, Fuchs L, Piel J, Von Mering C, Sourjik V, Hardt W-D. 2023. Chemotaxis and autoinducer-2 signalling mediate colonization and contribute to co-existence of Escherichia coli strains in the murine gut. Nat Microbiol 8:204–217.

19. Berne C, Brun YV. 2019. The Two Chemotaxis Clusters in Caulobacter crescentus Play Different Roles in Chemotaxis and Biofilm Regulation. Journal of Bacteriology 201:e00071–19.

20. Jenal U, Reinders A, Lori C. 2017. Cyclic di-GMP: second messenger extraordinaire. 5. Nat Rev Microbiol 15:271–284.

21. Hengge R. 2009. Principles of c-di-GMP signalling in bacteria. Nat Rev Microbiol 7:263– 273.

22. Tal R, Wong HC, Calhoon R, Gelfand D, Fear AL, Volman G, Mayer R, Ross P, Amikam D, Weinhouse H, Cohen A, Sapir S, Ohana P, Benziman M. 1998. Three cdg Operons Control Cellular Turnover of Cyclic Di-GMP in Acetobacter xylinum: Genetic Organization and Occurrence of Conserved Domains in Isoenzymes. Journal of Bacteriology 180:4416–4425.

23. Römling U, Galperin MY, Gomelsky M. 2013. Cyclic di-GMP: the First 25 Years of a Universal Bacterial Second Messenger. Microbiology and Molecular Biology Reviews 77:1– 52.

24. Khan F, Jeong G-J, Tabassum N, Kim Y-M. 2023. Functional diversity of c-di-GMP receptors in prokaryotic and eukaryotic systems. Cell Communication and Signaling 21:259.

25. Govers SK, Jacobs-Wagner C. 2020. Caulobacter crescentus: model system extraordinaire. Current Biology 30:R1151–R1158.

26. Bodenmiller D, Toh E, Brun YV. 2004. Development of Surface Adhesion in Caulobacter crescentus. J Bacteriol 186:1438–1447.

27. Aldridge P, Paul R, Goymer P, Rainey P, Jenal U. 2003. Role of the GGDEF regulator PleD in polar development of Caulobacter crescentus. Molecular Microbiology 47:1695–1708.

28. Wetmore KM, Price MN, Waters RJ, Lamson JS, He J, Hoover CA, Blow MJ, Bristow J, Butland G, Arkin AP, Deutschbauer A. 2015. Rapid quantification of mutant fitness in diverse bacteria by sequencing randomly bar-coded transposons. mBio 6:e00306–00315.

29. Hellenbrand CN, Stevenson DM, Gromek KA, Amador-Noguez D, Hershey DM. 2024. A deoxynucleoside triphosphate triphosphohydrolase promotes cell cycle progression in Caulobacter crescentus. bioRxiv 10.1101/2024.04.25.591158.

30. Jumper J, Evans R, Pritzel A, Green T, Figurnov M, Ronneberger O, Tunyasuvunakool K, Bates R, Žídek A, Potapenko A, Bridgland A, Meyer C, Kohl SAA, Ballard AJ, Cowie A, Romera-Paredes B, Nikolov S, Jain R, Adler J, Back T, Petersen S, Reiman D, Clancy E, Zielinski M, Steinegger M, Pacholska M, Berghammer T, Bodenstein S, Silver D, Vinyals O, Senior AW, Kavukcuoglu K, Kohli P, Hassabis D. 2021. Highly accurate protein structure prediction with AlphaFold. Nature 596:583–589.

31. Chan C, Paul R, Samoray D, Amiot NC, Giese B, Jenal U, Schirmer T. 2004. Structural basis of activity and allosteric control of diguanylate cyclase. Proceedings of the National Academy of Sciences 101:17084–17089.

32. Christen B, Christen M, Paul R, Schmid F, Folcher M, Jenoe P, Meuwly M, Jenal U. 2006. Allosteric Control of Cyclic di-GMP Signaling*. Journal of Biological Chemistry 281:32015– 32024.

33. Römling U, Liang Z-X, Dow JM. 2017. Progress in Understanding the Molecular Basis Underlying Functional Diversification of Cyclic Dinucleotide Turnover Proteins. J Bacteriol 199.

34. Lele PP, Hosu BG, Berg HC. 2013. Dynamics of mechanosensing in the bacterial flagellar motor. Proceedings of the National Academy of Sciences 110:11839–11844.

35. Tipping MJ, Delalez NJ, Lim R, Berry RM, Armitage JP. 2013. Load-Dependent Assembly of the Bacterial Flagellar Motor. mBio 4:10.1128/mbio.00551-13.

36. Antani JD, Sumali AX, Lele TP, Lele PP. 2021. Asymmetric random walks reveal that the chemotaxis network modulates flagellar rotational bias in Helicobacter pylori. eLife 10:e63936.

37. Laganenka L, Sourjik V. 2018. Autoinducer 2-Dependent Escherichia coli Biofilm Formation Is Enhanced in a Dual-Species Coculture. Applied and Environmental Microbiology 84:e02638–17.

38. Baker AE, Webster SS, Diepold A, Kuchma SL, Bordeleau E, Armitage JP, O’Toole GA. 2019. Flagellar Stators Stimulate c-di-GMP Production by Pseudomonas aeruginosa. J Bacteriol 201:e00741–18.

39. Newell PD, Monds RD, O’Toole GA. 2009. LapD is a bis-(3′,5′)-cyclic dimeric GMP-binding protein that regulates surface attachment by Pseudomonas fluorescens Pf0–1. Proc Natl Acad Sci U S A 106:3461–3466.

40. Dahlstrom KM, Giglio KM, Collins AJ, Sondermann H, O’Toole GA. 2015. Contribution of Physical Interactions to Signaling Specificity between a Diguanylate Cyclase and Its Effector. mBio 6:e01978–01915.

41. Dahlstrom KM, Giglio KM, Sondermann H, O’Toole GA. 2016. The Inhibitory Site of a Diguanylate Cyclase Is a Necessary Element for Interaction and Signaling with an Effector Protein. Journal of Bacteriology 198:1595–1603.

42. Giacalone D, Smith TJ, Collins AJ, Sondermann H, Koziol LJ, O’Toole GA. 2018. Ligand-Mediated Biofilm Formation via Enhanced Physical Interaction between a Diguanylate Cyclase and Its Receptor. mBio 9:10.1128/mbio.01254-18.

43. Flynn JM, Neher SB, Kim YI, Sauer RT, Baker TA. 2003. Proteomic discovery of cellular substrates of the ClpXP protease reveals five classes of ClpX-recognition signals. Mol Cell 11:671–683.

44. Tan W, Cheng S, Li Y, Li X-Y, Lu N, Sun J, Tang G, Yang Y, Cai K, Li X, Ou X, Gao X, Zhao G-P, Childers WS, Zhao W. 2022. Phase separation modulates the assembly and dynamics of a polarity-related scaffold-signaling hub. Nat Commun 13:7181.

45. Shin Y, Brangwynne CP. 2017. Liquid phase condensation in cell physiology and disease. Science 357:eaaf4382.

46. Saurabh S, Chong TN, Bayas C, Dahlberg PD, Cartwright HN, Moerner WE, Shapiro L. 2022. ATP-responsive biomolecular condensates tune bacterial kinase signaling. Science Advances 8:eabm6570.

47. Sobol G, Hershey DM. 2026. The gac system integrates physical and chemical cues to promote plant root attachment. Applied and Environmental Microbiology 92:e00778–26.

48. Ducret A, Quardokus EM, Brun YV. 2016. MicrobeJ, a tool for high throughput bacterial cell detection and quantitative analysis. 7. Nat Microbiol 1:1–7.

